# The heterogeneity of dermal mesenchymal cells reproduced in skin equivalents regulates barrier function and elasticity

**DOI:** 10.1101/2024.12.08.627431

**Authors:** Shun Kimura, Sachiko Sekiya, Sawa Yamashiro, Tetsutaro Kikuchi, Masatoshi Haga, Tatsuya Shimizu

## Abstract

The heterogeneity of dermal mesenchymal cells, including perivascular mesenchymal cells and papillary and reticular fibroblasts, plays critical roles in skin homeostasis. Herein, we present human skin equivalents (HSEs), in which pericytes, papillary fibroblasts, and reticular fibroblasts are spatially organized through autonomous three-cell interactions among epidermal keratinocytes, dermal fibroblasts, and vascular endothelial cells. The replication of dermal mesenchymal cell heterogeneity enhances skin functions, including epithelialization, epidermal barrier formation, and dermal elasticity, enabling *in vitro* evaluation of drug efficacy using methodologies that are identical to those used in human clinical studies. Furthermore, ascorbic acid-induced epidermal turnover and synthesis of well-aligned extracellular matrix via perivascular niche cells play crucial roles in improving skin barrier function and elasticity. Therefore, HSEs with heterogeneous dermal mesenchymal cells may improve our understanding of the mechanisms underlying skin homeostasis through cell-to-cell communication and serve as a model to animal experiments for developing precision medicine.

## Introduction

Intercellular crosstalk based on cellular heterogeneity plays a crucial role in maintaining the morphological and functional homeostasis of organs and has garnered attention as a potential target for studies on precision medicine^1^. Given that the integumentary organ system (IOS) is the most readily accessible organ system in the human body, the analysis of heterogeneity in human skin cells has established paradigms for scientific research in areas such as cell adhesion, inflammation, and tissue stem cells^2^. The IOS, which includes the skin, skin appendages, nerves, and blood vessels, plays essential roles in waterproofing, cushioning, protecting relatively deep tissues, excreting waste and thermoregulation^3^. Dermal mesenchymal cells contribute physical resilience by synthesizing extracellular matrix (ECM) components, such as collagen and elastin, and play a pivotal role in maintaining the homeostasis of epidermis, hair follicles, and vasculature structure via intercellular communication involving growth factors and cell adhesion^4,5^. Histological analyses and single-cell transcriptomes of the adult human skin have identified various subtypes of dermal mesenchymal cells, including papillary fibroblasts, reticular fibroblasts, pro-inflammatory fibroblasts, mesenchymal fibroblasts, dermal papilla cells, arrector pili muscle fibroblasts, pericytes, and mesenchymal stem cells^6–10^. The heterogeneity of dermal mesenchymal cells is modulated by factors, such as age, anatomical location, both intrinsic and extrinsic stressors, thereby influencing physiological processes including wound healing, fibrosis, atopic dermatitis, and aging^11–13^. Interestingly, the potential for intervention in aberrant alterations of heterogeneity of dermal mesenchymal cell is being increasingly demonstrated, as exemplified by the suppression of ultraviolet-induced selective depletion of papillary fibroblasts through local cyclooxygenase 2 inhibition, and attenuation of aging-associated transcriptomic changes in dermal fibroblasts by caloric restriction^14,15^. Therefore, elucidating the functional characteristics of dermal mesenchymal cell subtypes and mechanisms that maintain homeostasis of the heterogeneity may significantly contribute to developing precision medical technologies targeting skin diseases and aging.

The identification and determination of functional characteristics of dermal mesenchymal cell heterogeneity have been conducted through *in vivo* studies in mice and humans. However, animal experimentation requires careful consideration of interspecies differences with humans and faces increasing societal demands for a reduction^16–19^. In contrast, human clinical studies are valuable for basic research and for validating efficacy; however, they are not suitable for large-scale screening of functional materials. To enable societal implementation dermal mesenchymal cell heterogeneity, developing *in vitro* research models that can replicate and assess dermal cellular heterogeneity and organ-level functionality is necessary. Human cell-based *in vitro* research models such as two-dimensional (2D)-tissue cultured cells, skin organoids, and human skin equivalents (HSEs) are being explored as preclinical research tools for avoiding animal experimentation^20^. Although 2D tissue culture offers benefits for high-throughput screening, numerous subtypes of dermal mesenchymal cells, including papillary fibroblasts and pericytes, progressively lose their distinctive gene expression patterns and cellular morphology with successive passaging^7,21^. Spheroid culture-based organoids, which allow for sophisticated *in vitro* reconstruction of skin organ structures, including hair follicles, vasculature, and nerves, and the cellular heterogeneity of their constituent components, hold significant potential for advancing the understanding of cell–cell interactions and for future integration into drug screening platforms^22–25^. HSEs, distinguished by their capacity to fabricate tissues of customizable shapes at the centimeter scale, are already employed in the healthcare industry as commercially available platforms for tissue-level evaluation *in vitro*, encompassing drug safety, permeability, moisturizing barrier function, and mechanobiological aspects of skin physiology^26–28^.

Basic HSEs, including those commercially available, comprise a fully stratified epidermal layer, along with a dermal equivalent formed from three-dimensional (3D) ECM such as a decellularized dermis or a Type 1 collagen seeded with dermal fibroblasts ^29,30^. The advancement of HSEs that relatively closely replicate the structure and functionality of native human skin is actively progressing by incorporating additional cell types, such as fibroblast subpopulations, vascular cells, immune cells, neural cells, and adipocytes, along with the integration of spheroid cultured tissues and application of 3D bioprinting technologies^31–33^. The vascular network, which is indispensable for delivering oxygen and nutrients and regulating of inflammatory responses, has been integrated into HSEs by incorporating vascular endothelial cells within the dermal compartment, transplanting endothelial cells into bioprinted vasculature-like structures, and implementing perfusion culture systems^34,35^. These innovations have facilitated the application of HSEs in drug delivery research^36^. Although the development of HSEs that reconstruct the cellular heterogeneity of native skin remains relatively limited, recent single-cell transcriptomic analyses have demonstrated that HSEs recapitulate the principal *in vivo* epidermal keratinocyte subtypes, and encompass unique keratinocyte populations exhibiting aberrant signaling pathways associated with epithelial– mesenchymal transition. Intriguingly, the appearance of these artificial cell subtypes is rescued by incorporating fibroblasts into dermal equivalents and xenografting onto immunodeficient mice, indicating that intercellular communication with keratinocytes and dermal mesenchymal cells plays a crucial role in maintaining the homeostasis of epidermal cell heterogeneity^37^. In contrast, the heterogeneity of dermal mesenchymal cells of HSEs remains largely unexplored. To investigate the functional roles of distinct dermal mesenchymal cell subpopulation, HSEs have been constructed using papillary fibroblasts and pericytes derived from primary cultures of human dermis. These models have revealed that epithelialization is strongly influenced by intercellular communication, including Wnt signaling and laminin subunit alpha 5 (LAMA5)-mediated pathways^7,21,38,39^. However, in constructing HSEs that replicate the heterogeneity of dermal mesenchymal cells, the strategy for isolating, culturing, and spatially arranging each cellular subpopulation is economically impractical.

In developing and regenerating different organs, cell-to-cell communication plays an essential role in inducing and maintaining differentiation of constituent cells19. A specific subset of tissue-resident mesenchymal cells located close to blood vessels plays a key role in epidermal differentiation and ECM production4,40,41. Therefore, we hypothesized that by vascularizing an HSE and reproducing cell-to-cell communication on the basis of reconstructed perivascular niche, constructing an HSE that relatively faithfully replicates the *in vivo* characteristics of skin would be possible. In this study, we report an HSE that replicates the heterogeneity of human dermal mesenchymal cells, including pericytes, papillary fibroblasts, and reticular fibroblasts, by utilizing commercially available primary cells, specifically normal human epidermal keratinocytes (NHEKs), normal human dermal fibroblasts (NHDFs), and human umbilical vein endothelial cells (HUVECs). Additionally, we demonstrate that our HSEs serve as a valuable model for elucidating the role of dermal mesenchymal cell heterogeneity in preserving skin structural integrity and functional homeostasis.

## Results

### Tricellular communication among epidermal keratinocytes, dermal fibroblasts, and vascular endothelial cells enhances the organization of skin and blood vessels

To investigate the role of communication among epidermal keratinocytes, dermal fibroblasts, and vascular endothelial cells in skin tissue formation, we reconstructed seven types of HSEs, each incorporating one to three types of these constituent cells. The HSEs were named using the initials of the incorporated cell types. Specifically, HSEs composed of a single cell type, epidermal keratinocytes, dermal fibroblasts, or vascular endothelial cells, were referred to as the E, D, or V models, respectively. Models containing two of these cell types were designated as the DV, EV, and ED models, and the model incorporating all three cell types was referred to as the EDV model (Fig. 1a and 1b). All HSEs were constructed using a previously reported method for reproducing tensional homeostasis in a skin model^27^. After 14 d of air lift culture, the reconstructed tissue was 12 mm in diameter and 1–2 mm in thickness. The E and EV models were transparent and extremely soft (Fig. 1b). Hematoxylin and eosin (H&E) staining was used to visualize the regulation of skin and blood vessel organization through cell-to-cell communication (Fig. 1c). In the E and EV models, an abnormal epidermis lacking a basal cell layer formed, whereas normal epithelialization was observed in the ED and EDV models. This finding indicates that crosstalk between NHEKs and NHDFs is essential for epithelialization. Endothelial tubes in the dermis were observed only sparsely in the V and EV models; however, they were distributed throughout the skin in the DV and EDV models. In particular, the endothelial tubes in the EDV model tended to orient horizontally along the skin surface and increased in number with increasing distance from the epidermis. Sirius red staining revealed localized collagen fiber deposition (Fig. 1d). In the ED model, collagen fiber formation was confirmed at the epidermal–dermal junction because of the well-known interaction between the epidermis and dermis. In the DV and EDV models, the distribution of collagen deposition within the dermis was similar to that of the endothelial tubes. Interestingly, collagen deposition at the epidermal–dermal junction and around endothelial tubes was promoted in the EDV model compared to that in the ED and DV models.

**Figure 1.**
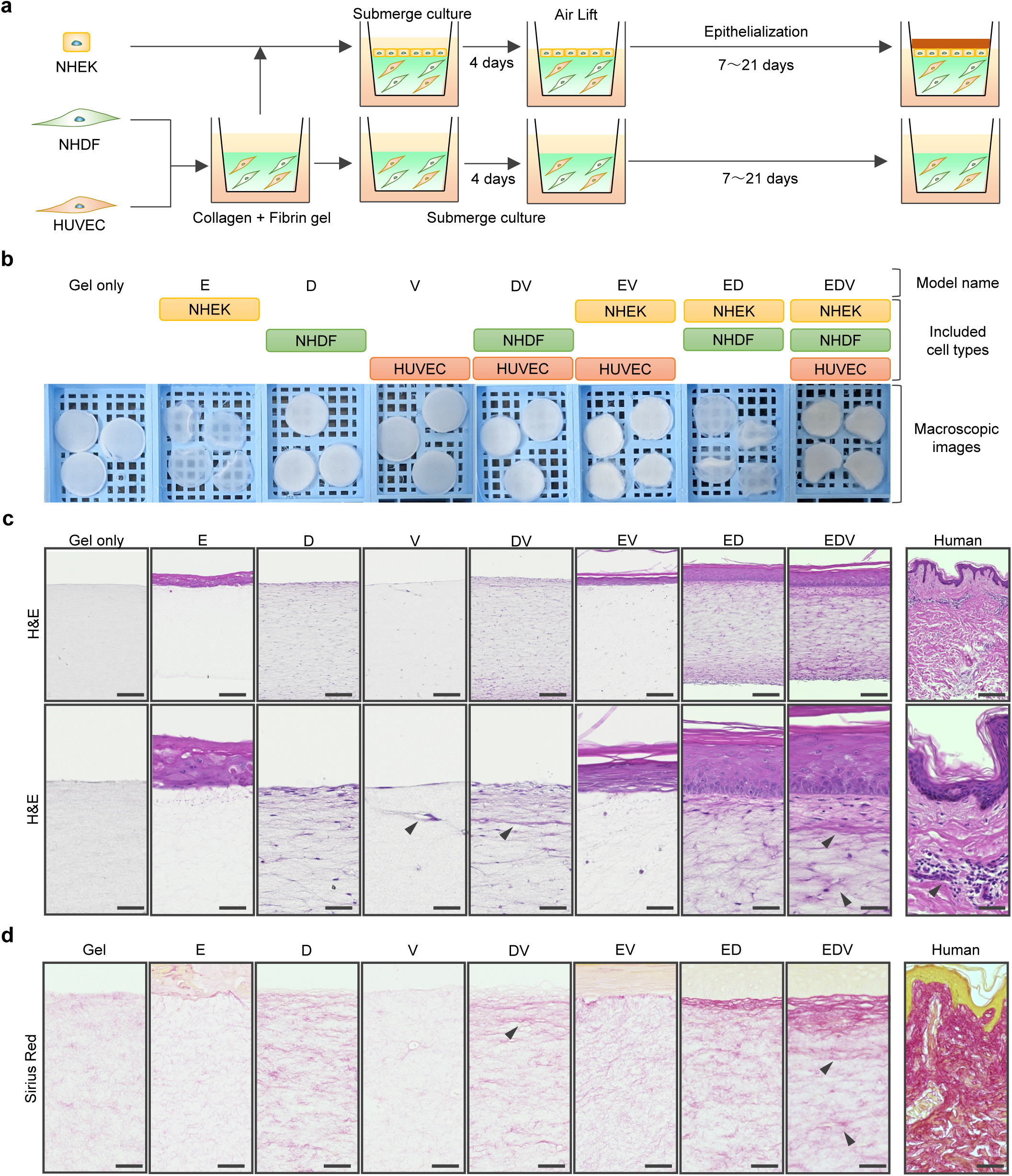
Tricellular communication regulates skin and vascular organization. (a) Schematic of the construction method of HSEs. (b) Macroscopic images of the HSEs. Scale bar, 1 mm. (c) H&E-stained images of human skin and the HSEs at low (upper columns; scale bar, 200 µm) and high (lower columns; scale bar, 50 µm) magnification. The arrowheads indicate the endothelial tube. (d) Sirius red-stained images of human skin and the HSEs; scale bar, 50 µm.

The morphogenesis of the epidermis, endothelial tube, and dermis in each HSE was analyzed by using immunohistology. The results of epidermal marker staining demonstrated that the epidermis of the ED and EDV models exhibited a cytokeratin 5 (CK5)-positive basal layer, a claudin 1 (CLDN1)-positive basal to granular layer, filaggrin (FLG)-positive keratohyalin granules, and a cutokeratin 10 (CK10)-positive stratum corneum, closely resembling the epidermis of natural human skin (Fig. 2a). Furthermore, the number of Ki67-positive cells was highest in the EDV model and relatively low in the ED, EV, and E models (Fig. 2b). Trans-epidermal water loss (TEWL) in the EDV model was 9.68 g/m^2^h, and water evaporation was significantly suppressed compared to that in the ED model (Fig. 2c). These results indicate that dermal fibroblast–epidermal keratinocyte communication, that promotes epithelialization and barrier function is enhanced by vascular endothelial cells.

**Figure 2.**
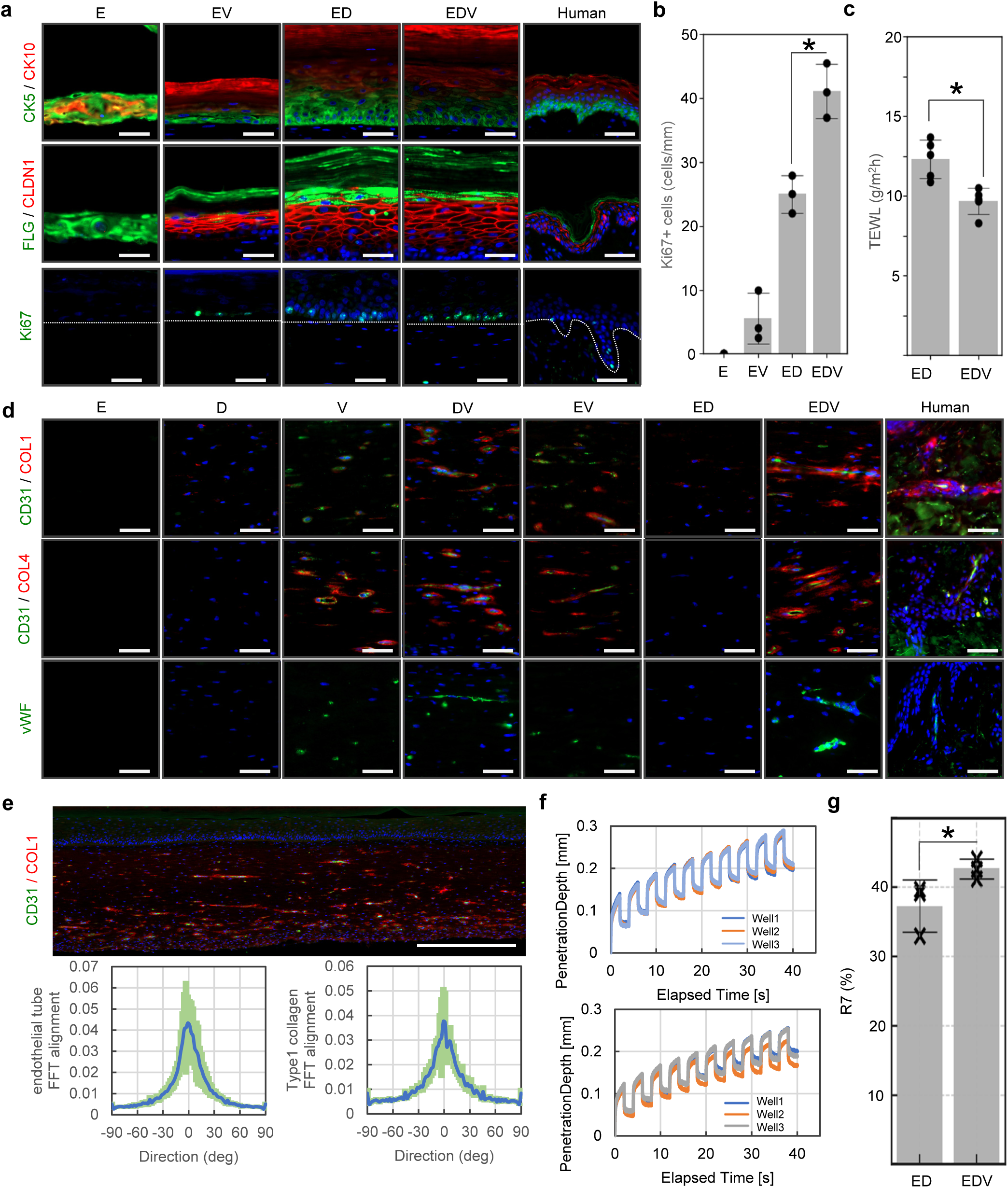
Tricellular communication enhances skin barrier function and elasticity by inducing self-organization. (a) Immunohistochemical analyses of human skin and the HSEs for evaluating epithelialization. The dotted lines indicate the boundary between the epidermis and dermis. Scale bar, 50 µm. (b) Analysis of the Ki67-positive basal keratinocyte ratio in HSEs. *P < 0.001 (Dunnett’s test following two-way analysis of variance). (c) TEWL analysis of the ED and EDV models *P < 0.05 (two-tailed Student’s t–tests; error bars represent standard deviation). (d) Immunohistochemical analyses of human skin and HSEs for evaluating angiogenesis and dermal collagen synthesis. (e) Macroscopic image of CD31 and COL1 distribution (upper column) and FFT analysis (lower columns) of the alignment of CD31-positive endothelial tubes and COL1-positive type 1 collagen fibers in the EDV model. Scale bar, 500 µm. (f) Skin deformation curves of the ED (upper panel) and EDV models (lower panel). (g) Evaluation of R7 factors of the ED and EDV models. *P < 0.05 (two-tailed Student’s t-tests; error bars represent standard deviation).

Immunostaining for the endothelial cell markers CD31 and von Willebrand factor (vWF) revealed active endothelial tube formation in the presence of both HUVECs and NHDFs (Fig. 2d). In the DV and EDV models, deposition of collagen types 1 and 4 was observed around the endothelial tubes, suggesting fibroblast– endothelial cell crosstalk stabilizes the endothelial tube structure through ECM synthesis (Fig. 2d). In the ED model, formation of type 1 collagen fiber in the dermis was stimulated around the epidermal–dermal junction, whereas in the EDV model, additional strong type 1 collagen expression originating from endothelial tubes was observed, resulting in the formation of collagen fibers that spread throughout the entire dermis. Microcirculation in natural human skin is organized into horizontal plexuses situated 1–1.5 mm below the skin surface^40^. Immunostaining for CD31 and type 1 collagen revealed that the blood vessels and collagen fiber in the EDV model were distributed in parallel to the epidermal plane, and mathematical assessment using 2D fast Fourier transform (2D-FFT) analysis corroborated this observation (Fig. 2e). The collagen network, which is arranged parallel to the skin surface, functions to withstand mechanical forces acting perpendicular to the skin. Therefore, we evaluated the elasticity of HSEs, a physical indicator of skin aging and correlating factor of wrinkles and sagging, using the Cutometer-based protocol applied in human clinical studies^41^. Skin elasticity could be measured only in the ED and EDV models because the tissue in the other HSEs was damaged by negative pressure loading. The ED and EDV models withstood 10 negative pressure loading cycles (Fig. 2f). The R7 value, which is an elasticity index reportedly associated with facial sagging, increased in the EDV model (Fig. 2g).

Therefore, tricellular interactions among epidermal keratinocytes, dermal fibroblasts, and endothelial cells play critical roles in epidermal turnover, collagen synthesis, and angiogenesis and are sufficient to enhance tissue-scale skin functionality, including viscoelastic properties, and integrity of the epidermal barrier.

### Cell-to-cell communication in a 3D environment replicates dermal mesenchymal cell heterogeneity that is lost in 2D culture

To evaluate the impact of vascularization on fibroblast heterogeneity in the HSEs, we analyzed the spatial distribution of cells with positive markers for papillary fibroblasts (CD39 and fibroblast activation protein [FAP]), reticular fibroblasts (CD36 and CD90), and pericytes neural/glial antigen (NG2) and alpha-smooth muscle actin (αSMA) via immunohistochemistry (Fig. 3a and 3b). Fibroblasts in the D model were positive for the pan-fibroblast marker vimentin; however, the expression of all markers in papillary and reticular fibroblasts and pericytes was barely detectable. CD39-positive cells were observed in the DV and EDV models, and FAP-positive cells were observed in the ED and EDV models. Interestingly, CD39-positive cells tended to localize around the endothelial tubes, whereas FAP-positive cells were observed mainly near the epidermal– dermal junction; similar patterns were also observed in natural human skin. CD36-positive cells were distributed in the DV and EDV models, and CD90-positive cells were sparsely distributed in the ED, and EDV models (Fig. 3a). NG2- and αSMA-positive pericytes were localized to the periphery of the lumen formed by vWF-positive vascular endothelial cells (Fig. 3b).

**Figure 3.**
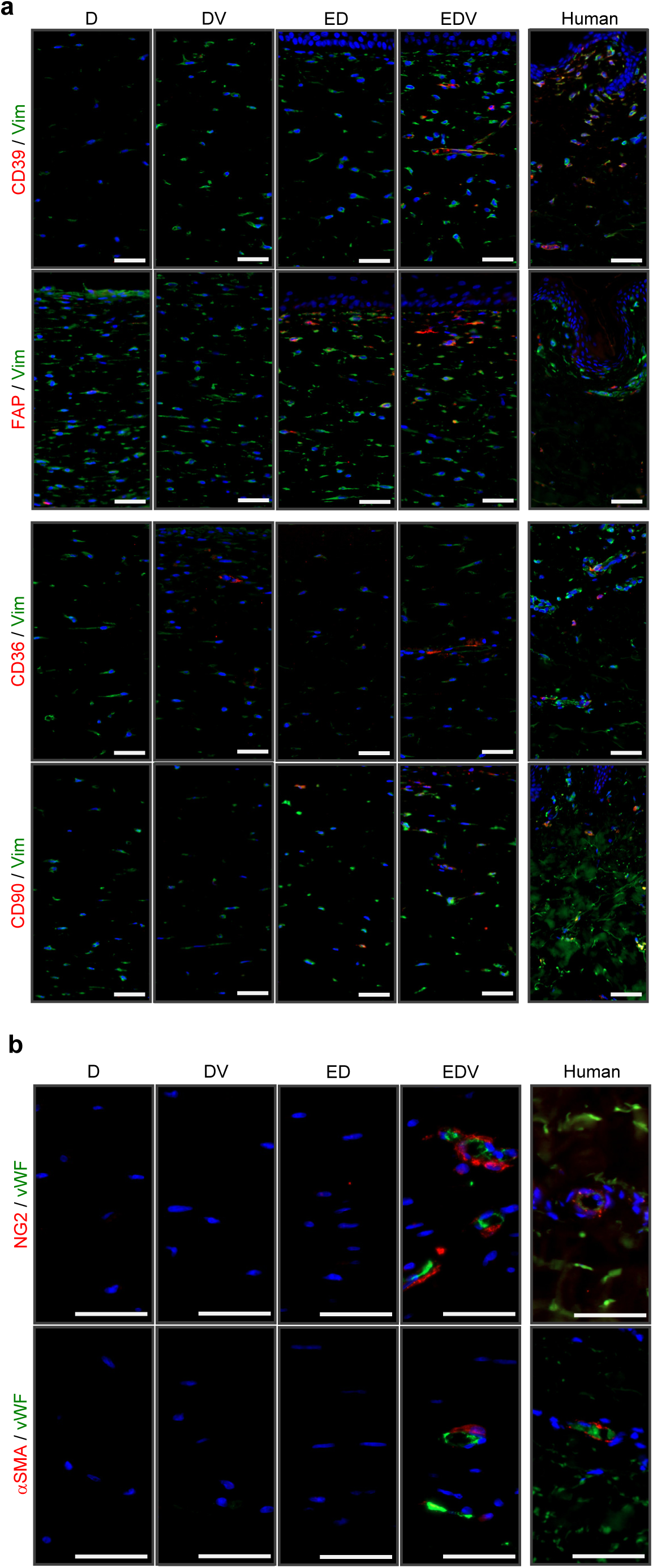
Immunohistological analysis of dermal mesenchymal cell distribution in the HSEs and human skin. (a) Immunohistochemical analyses of ED, EDV and human skin samples to evaluate the presence and distribution of dermal mesenchymal cell subpopulations. Scale bar, 50 µm. (b) Immunohistochemical analysis of pericyte distribution in ED, EDV, and human skin tissues. Scale bar, 50 µm.

Immunohistochemical analysis revealed that the fibroblasts constituting the HSEs exhibited heterogeneity in terms of expression of the cell surface markers, depending on the constituent cell types. Therefore, we conducted single-cell RNA sequencing (scRNA-seq) to analyze the single-cell transcriptomes of dermal mesenchymal cell populations within the HSEs. We profiled three samples: the ED and EDV models cultured at the air–liquid interface for 14 d, a mixture of NHEKs, NHDFs, and HUVECs cultured separately under 2D conditions (referred to as 2D) (Fig. 4a). In an initial analysis, we obtained an overview of diverse cell populations by integrating cells from three samples. After quality filtering, a total of 23,911 cell profiles were successfully obtained across all conditions. A uniform manifold approximation and projection (UMAP) plot revealed three major cell populations and 20 distinct clusters (Fig. 4b, left). Based on the existing marker genes, we identified four cell types (keratinocytes, fibroblasts, vascular endothelial cells, and pericytes) for each cell culture (Fig. 4b, right). The UMAP plot illustrates specific impact of each culture condition on fibroblast population (Supplementary Fig. 1a). The major fibroblast clusters under 2D, #1, #6, #13, and #17 were relatively minor subpopulations in the ED and EDV models. The clusters #0, #2, #9, #10, and #11 were major in both the ED and EDV models; however, cluster #14, characterized by the expression of pericyte marker genes (*ACTA2* and *RGS5*), was detected as a subpopulation unique to the EDV model (Fig. 4c). We hypothesized that the formation and maintenance of cluster #14 involves endothelial cell–fibroblast communication and investigated this interaction using CellChat analysis^42^. Cluster #14 showed increased intracellular communication with vascular endothelial cell clusters through vascular endothelial growth factor (VEGF)B–VEGF receptor 1 (VEGFR1), placental growth factr (PGF)–VEGFR1, laminin–dystroglycan, and laminin–integrin interactions compared to that by other fibroblasts clusters (Fig. 4d). The ED and EDV models promoted the expression of collagen fiber-related genes such as *COL1A1*, *COL3A1*, *COL4A1*, *COL4A2*, and *COL5A3*, but suppressed the expression of elastic fiber-related genes, including *ELN* and *FBN1*, compared to that in the 2D conditions (Fig. 4e). This change in the transcriptome was relatively highly pronounced in the EDV model, suggesting the contribution of cluster #14.

**Figure 4.**
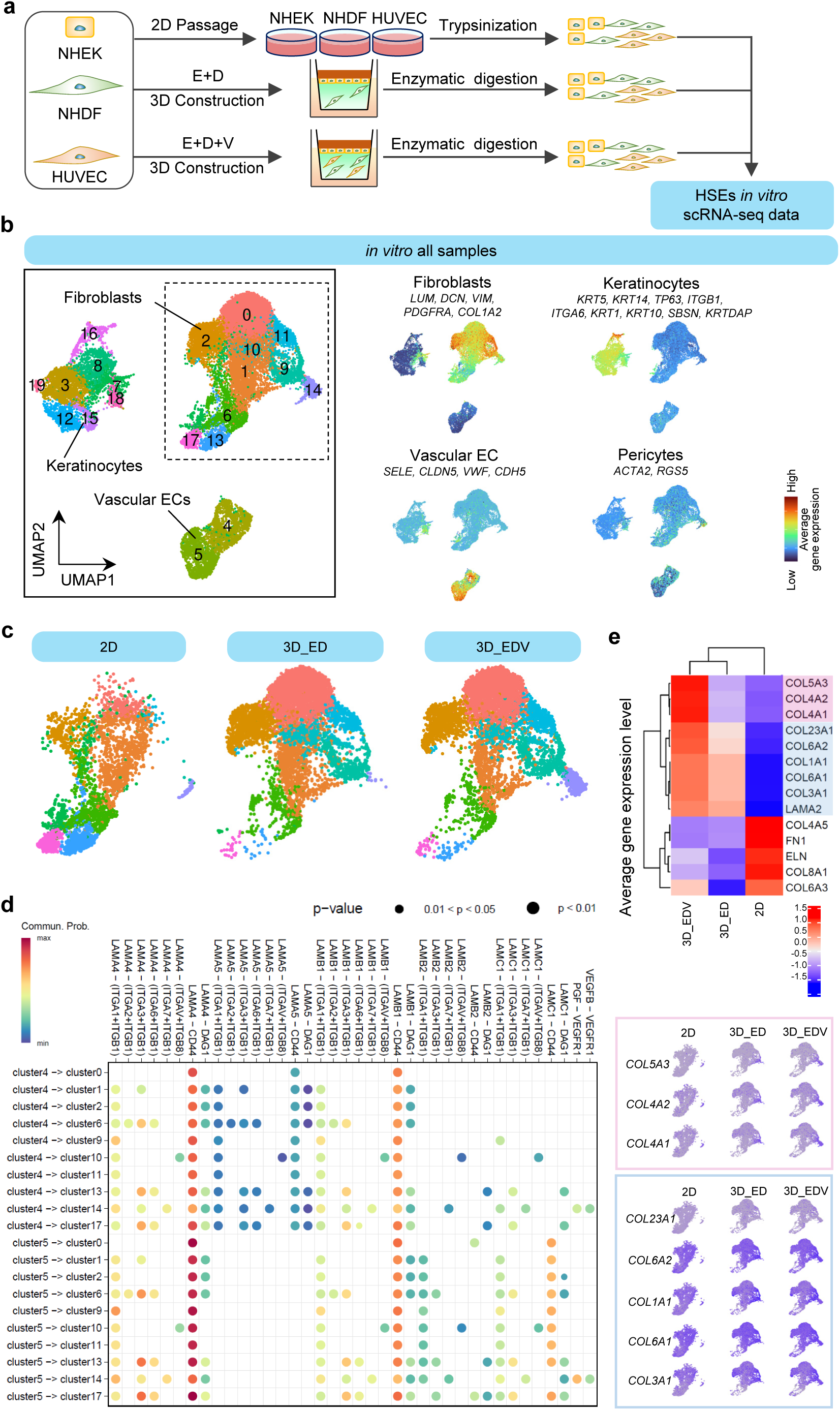
The 3D culture environment and cell-to-cell communication replicate dermal mesenchymal cell heterogeneity *in vitro*. (a) Illustration of the workflow for sampling and analysis of the scRNA-seq data. (b) UMAP plot of the scRNA-seq data for in *vitro* HSEs; the average expression of four cell-type markers projected onto the UMAP plot for identifing cell populations. (c) Distribution of fibroblasts and pericytes under each of the three conditions. (d) Cellular communication from vascular endothelial cells (ECs; Clusters #4 and #5) to pericytes in Cluster #14 and fibroblasts in Cluster #9. (e) Heatmap showing the average expression of ECM-related genes across all fibroblasts under different conditions.

To investigate the biological significance of dermal mesenchymal cell heterogeneity of the HSEs, we referred to a published *in vivo* human skin scRNA-seq dataset and integrated it with our HSE datasets using the R Harmony data integration algorithm^11^. A UMAP plot revealed four major cell populations and 11 distinct clusters (Fig. 5a). To distinguish the results of this integrated analysis from previous *in vitro* data analysis, apostrophes were added to the cluster numbers in this analysis. The four major clusters were defined as keratinocytes, fibroblasts, vascular endothelial cells, and pericytes according to the expression patterns of marker genes (Fig. 5b). In the ED and EDV models, similar to *in vivo* skin, fibroblasts were composed of mainly clusters #0’, #1’, and #8’. In contrast, 2D-cultured fibroblasts contained very few cells from clusters #0’ and #8’, whereas clusters #2’ and #11’, which are rarely found in natural human skin, comprised their main subpopulations (Fig. 5c). Cell cycle scoring using the Seurat scRNA-seq analysis pipeline (ccSeurat)^43^ (https://satijalab.org/seurat/v3.1/cell_cycle_vignette.html) revealed that cells in cluster #2’, characteristic of 2D-cultured fibroblasts, were in the G2/M and S phases (Fig. 5d). Four main fibroblast subpopulations that could be spatially localized and showed differential secretory, mesenchymal and proinflammatory functional annotations have been previously defined^11^. We reanalyzed this dataset for identifying these four fibroblast subtypes and pericytes (Supplementary Fig. 2). We then plotted the cells corresponding to each subtype on the integrated UMAP plot (Fig. 5e). Next, we analyzed the clustering ratios within the integrated UMAP plot for each of the five natural human skin subpopulations of cells including secretory papillary, secretory reticular, proinflammatory, mesenchymal fibroblasts and pericytes (Fig. 5f). The results revealed that secretory papillary and pro-inflammatory fibroblasts corresponded to cluster #0’; secretory reticular fibroblasts corresponded to cluster #1’; mesenchymal fibroblasts corresponded to cluster #8’; and pericytes corresponded to cluster #6’. We also investigated the Gene Ontology (GO) terms enriched in cluster #0’; that represents the papillary fibroblast phenotype and cluster #1’; that represents the reticular fibroblast phenotype. In cluster #1’; GO terms related to collagen and elastic fiber formation were particularly enriched, suggesting that cells in cluster #1’ exhibited a phenotype closely resembling that of reticular fibroblasts (Fig. 5g).

**Figure 5.**
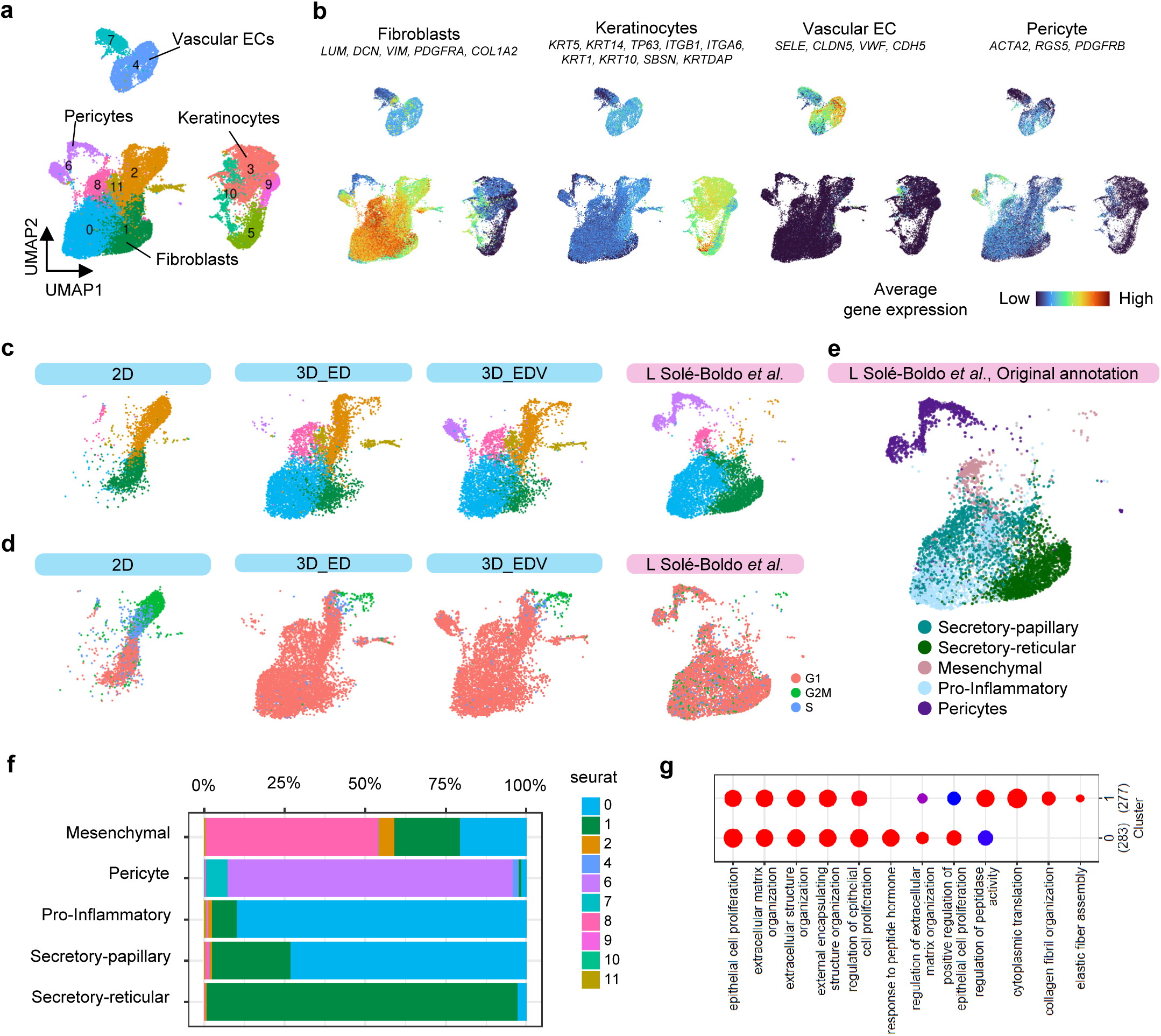
Biological significance of fibroblast heterogeneity in HSEs through integrated analysis of human scRNA-seq data. (a) UMAP plot of the scRNA-seq data for integrating HSEs and *in vivo* skin data. (b) Average expression of established cell type markers was projected on the UMAP plot to identify cell populations. Red indicates maximum gene expression, whereas blue indicates low or no expression of a particular set of genes in log-normalized UMI counts. (c) UMAP plot integrating the scRNA-seq data of HSEs and *in vivo* skin, showing the distribution of fibroblasts and pericytes across the three *in vitro* and *in vivo* conditions. (d) Cell cycle status of fibroblasts under each condition. (e) UMAP plot showing the correspondence of the four fibroblast subpopulations and pericytes, as previously reported^11^. (f) Proportion of cells from each subpopulation as previously defined^11^ and their clustering into specific clusters, as shown in Fig. 4e. (g) Enriched GO terms in clusters #0’ and #1’.

### Nutrient-poor culture conditions in the HSEs induce a senescent**–**like phenotype, which is rescued by ascorbic acid

Since blood vessels are central to maintaining organ homeostasis, vascular aging is hypothesized to be a fundamental upstream factor of organismal aging^44,45^. During skin aging, a reduction in blood vessels within the papillary layer and decline in their functions are thought to contribute to thinning of the epidermis and papillary dermis owing to impaired supply of nutrient and paracrine factors from vascular endothelial cells; however, experimental evidence for this phenomenon remains limited^46–48^. Therefore, to evaluate the impact of plasma-derived nutrient deprivation induced by peripheral vascular reduction, ED and EDV were cultured for 10 d in a nutrient poor (NP) medium characterized by low FBS concentration and lack of basic fibroblast growth factor (bFGF) and ascorbic acid (AA). Moreover, ascorbic acid, a well-evaluated antiaging material known for its antioxidant properties, ability to induce epithelialization, and role as a coenzyme in collagen synthesis^49–51^, was added to the NP medium for evaluating its impact on HSE organization and skin functionality. H&E staining and immunohistological analysis revealed that skin tissue, including the epidermis, dermis, and endothelial tubes with pericytes, was reconstructed even under NP culture conditions (Fig. 6a and 6b). However, the number of Ki67-positive cells in the epidermal basal cell layer was significantly lower in the EDV model than in the ED model (Fig. 6c). When 500 μM AA was added, thickening of the CK5- and/or CK10-positive epidermal layer and promotion of type 1 collagen deposition were observed in both the ED and EDV models (Fig. 6b). AA increased the number of Ki67-positive cells in both the ED and EDV models; however, the EDV model showed a significantly relatively high number in the number of Ki67-positive cells (Fig. 6c). In both the ED and EDV models, ascorbic acid reduced TEWL and enhanced skin barrier function (Fig. 6d). A 2D-FFT analysis revealed that, NP culture conditions with AA resulted in horizontal orientation of CD31-positive vascular endothelial cells and the surrounding type 1 collagen relative to the epidermal layer, compared to that of the NP alone culture conditions (Fig. 6e). Furthermore, in the AA-depleted EDV model, collagen fragments of types 3 and 4 were deposited around CD31-positive endothelial tubes, suggesting collagen degradation (Fig. 6b). NG2- and αSMA-positive pericytes were observed surrounding the endothelial tubes regardless of the presence or absence of AA (Fig. 6b). Although no changes were detected in the ED model, skin elasticity was significantly increased by AA in the EDV model (Fig. 6f). In summary, NP culture conditions enabled *in vitro* replication of epidermal thinning, barrier disruption, reduction in dermal elasticity with imbalanced collagen synthesis and degradation, and disordered vascular orientation, characteristics similar to those of human skin aging. AA was effective in mitigating these aging-related changes. These findings demonstrate that our HSEs serve as a valuable tool for evaluating the efficacy of functional molecules, and suggest that vascular endothelial cells and pericytes play critical roles in the underlying mechanisms of action of these molecules.

**Figure 6.**
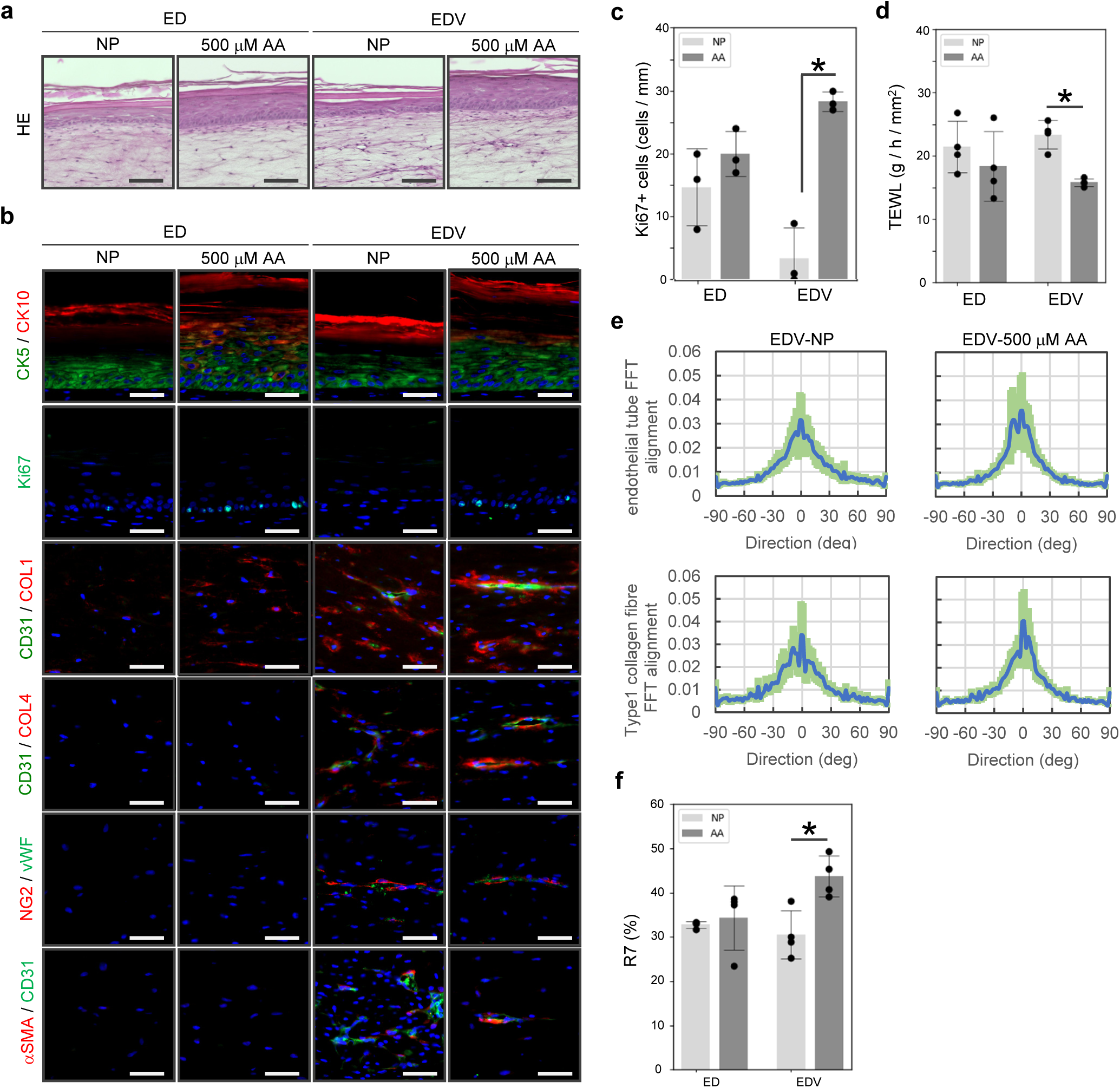
NP culture conditions in the HSEs induce a senescent phenotype, which is suppressed by AA. (a) H&E-stained images of skin tissue organization induced by AA supplementation in the HSEs cultured under NP conditions. Scale bar, 100 µm. (b) Immunohistochemical analysis of the HSEs for visualizing the impact of NP conditions and ascorbic acid on re-epithelialization, dermal ECM synthesis, and angiogenesis. Scale bar, 50 µm. (c) Analysis of the Ki67-positive basal keratinocyte ratio in the HSEs. *P < 0.001 (two-tailed Student’s t-tests; error bars represent standard deviation). (d) TEWL analysis of the ED and EDV models. *P < 0.001 (two-tailed Student’s t-tests; error bars represent standard deviation). (e) A 2D-FFT analysis (lower columns) of the alignment of CD31-positive endothelial tubes and COL1-positive type 1 collagen fibers in the EDV model. (f) Evaluation of R7 factors of the ED and EDV models. *P < 0.01 (two-tailed Student’s t-tests; error bars represent standard deviation).

## Discussion

In this study, we demonstrated that the heterogeneity of dermal mesenchymal cells, such as papillary and reticular fibroblasts and pericytes, in natural human skin can be partially reproduced by reconstructing an HSE using NHEKs, NHDFs, and HUVECs. In particular, the EDV model containing a pericyte-like subpopulation significantly improved skin and vascular tissue structure, barrier function, and elasticity, enabling functional evaluation according to human clinical protocols. These results elucidate the role of vascular cells in the developing the heterogeneity of dermal mesenchymal cell and demonstrate that the incorporation of vasculature into HSEs is an effective approach for recapitulating nutrient supply and inflammatory responses, as traditionally considered, and achieving a relatively highly advanced reconstruction of the skin transcriptome and functional properties. Our HSE, which reproduces fibroblast heterogeneity using stably available primary culture cells, provides a model alternative to animal experiments, which can be used to analyze the mechanisms controlling skin physiological functions based on cell-to-cell communication.

Papillary fibroblasts play important roles in wound healing functions such as re-epithelialization and collagen synthesis. Furthermore, papillary fibroblasts derived from elderly donors have reduced ability of proliferation and low secretion levels of keratinocyte growth factor, suggesting that this effect may involve loss of the papillary dermis, and elasticity, and deformation of skin texture owing to aging^38,39^. In fact, recent scRNA-seq analyses of human skin has revealed changes in fibroblast subpopulations in the skin owing to aging and disease, suggesting the possibility of developing new therapeutic strategies by controlling fibroblast heterogeneity^6,11,52^. Epidermal Wnt signals are involved in inducing differentiation of the papillary fibroblast lineage, and intervention in dermal fibroblast subpopulations controls skin morphogenesis and functionality in animals^53^. However, owing to the lack of appropriate *in vitro* models and ethical restrictions to clinical *in vivo* studies, the prospects of applying *in vitro* findings of human fibroblast heterogeneity to *in vivo* health care are unclear. In our HSE, fibroblasts induced by 3D culture were in the G1 phase and differentiated into papillary, reticular, and mesenchymal fibroblast-like populations, exhibiting properties similar to those of native skin fibroblasts. Interestingly, cells positive for FAP, a marker for papillary fibroblasts, were present near the epidermal–dermal junction in both the ED and EDV models; however, CD39-positive cells were observed only in the EDV model. This observation suggests that the keratinocyte–fibroblast communication is required for the induction and maintenance of papillary fibroblast differentiation, and that some subpopulations, such as CD39-positive cells, may require interaction with vascular endothelial cells. By elucidating the mechanisms controlling human fibroblast heterogeneity via cell tracking and spatiotemporal gene expression control techniques, our HSE may contribute to developing cell therapies for wound healing and aging.

In addition to papillary and reticular fibroblasts, perivascular mesenchymal cells, such as dermal stem cells and pericytes, play important roles in skin morphogenesis and function. Pericytes play an important role as the major source of ECM production during tissue regeneration in severl organs^54,55^. HSEs reconstructed using human pericytes and fibroblasts separated by flow cytometry have been reported to promote the proliferation of epidermal basal cells through LAMA5 expression by pericytes^21^. In our EDV model, epithelialization and dermal ECM synthesis were promoted with the emergence of a pericyte-like population. However, our HSE is unique in that it does not require fresh human skin and allows the construction of a model containing NG2- and αSMA-positive pericytes in a relatively simple process using commercially available subcultured cells. Pericytes are a heterogeneous cell population *in vivo*, and pericytes from the same tissue may originate from multiple sources^56,57^. Integrative analysis of scRNA-seq data from natural human skin and our HSEs revealed that although the pericyte-like population in HSEs clustered with human pericytes within the same subpopulation, their distributions on the UMAP plot only partially overlapped. At the very least, the induced pericyte-like cluster #14 replicated well-known *in vivo* angiogenic processes through interactions with endothelial cells, such as VEGFB and PGF signaling via VEGFR1, and basement membrane-mediated interactions between laminins and integrins^58–60^. These results suggest that the pericytes in our HSE partially replicate certain populations and functions of human pericytes, although the origin of this cell lineage and mechanisms driving their induction remain unclear. Cell tracking experiments using fluorescent dyes or genetic modifications targeting HSE constituent cells may contribute to addressing these gaps.

In addition to replicating the heterogeneity of dermal mesenchymal cells, the EDV model significantly improved dermal elasticity and skin barrier function. This improvement in dermal elasticity is attributed to the functionality of active ECM synthesis by pericytes, compared to that by other subpopulations, and the autonomous recapitulation of oriented collagen fiber formation along endothelial tubes. Furthermore, culturing under NP conditions enabled the HSE model to replicate the phenotype of aged skin, including epidermal barrier dysfunction owing to reduced proliferation of basal keratinocytes, diminished dermal elasticity owing to decreased collagen synthesis, increased collagen degradation by matrix metalloproteinases, and loss of vascular orientation. In natural human skin, oxidative stress and dermal ECM degradation cause vascular atrophy and reduces vascular density, leading to depletion of nutrient and oxygen supplies to the papillary dermis and epidermis, which contributes to aging. Our findings support this hypothesis. In our NP HSEs, a reduction in vascular density was not observed, which is likely attributed to VEGF in the medium continuously inducing angiogenesis. AA, which functions as both an antioxidant and a coenzyme for 2-oxoglutarate-dependent dioxygenases, is an effective skin–antiaging agent. However, the effects of AA on the vascular endothelium and perivascular mesenchymal cells and its impact on skin mechanical properties remain unclear. Our findings revealed that ascorbic acid increased collagen synthesis in both the ED and EDV models but significantly improved skin viscoelasticity only in the EDV model, indicating that vascular endothelial cells and pericytes play crucial roles in regulating skin elasticity and serve as target cells for AA. In summary, our model emphasizes the importance of vascular cells in the study of skin aging.

In conclusion, we reconstructed HSEs that reproduced skin fibroblast heterogeneity, tissue structure, and physiological function in natural human skin. Further studies, such as elucidating the mechanisms controlling fibroblast heterogeneity, evaluating the impact of each cell population on skin morphogenesis, and identifying drugs that can control the decrease or increase in specific fibroblast populations, may provide new insights into the fields of basic science, personalized medicine, and health care. Our model may help avoid animal experiments and contribute to the sustainable growth of the health care industry as a versatile tool for evaluating skin functionality as an alternative to clinical research. HSEs that highly replicate skin cell heterogeneity and function may be developed by improving their vascular structure through perfusion culture and providing cell-to-cell communication via organoids such as the IOS and nervous systems.

## Methods

### Cell culture

Normal human epidermal keratinocytes (NHEKs) were purchased from KURABO Industries Ltd. (Osaka, Japan) and were maintained in HuMedia-KG2 (KURABO). Normal human dermal fibroblasts (NHDFs) were purchased from KURABO and maintained in Dulbecco’s modified Eagle’s medium (Thermo Fisher Scientific, MA, USA) supplemented with 10% fetal bovine serum (Corning, NY, USA), 100 units ml^-1^ penicillin, and 100 µg ml^-1^ streptomycin (Thermo Fisher Scientific). Human umbilical vein endothelial cells (HUVECs) were purchased from KURABO and maintained in HuMedia-EG2 (KURABO). All cells were seeded into 150-mm dishes and grown in a humidified atmosphere of 5% CO_2_ at 37 °C.

### Reconstruction of the tension homeostatic skin (THS) model

The THS model was generated via the following procedure. A total of 24 × 10^5^ NHDFs and 30 × 10^5^ HUVECs were resuspended in 2.4 ml of 1.25 mg ml^-1^ bovine dermis-derived native collagen solution (KOKEN CO. Ltd., Tokyo, Japan) that was mixed with 1× DMEM (Thermo Fisher Scientific), 15 mM HEPES (Dojindo, Kumamoto, Japan), 10 mM NaHCO_3_ (FUJIFILM Wako Pure Chemical Corporation), and 5 mg ml^-1^ fibrinogen from bovine plasma type I-S (Merck KGaA, Darmstadt, Germany); then, the cells were plated in a high-density translucent-membrane cell culture insert (0.4-µm pore size) (Corning, NY, USA) in 6-well plates. After solidification, the dermal equivalents were held in place by a Snapwell culture insert (Corning). NHEKs were seeded at a density of 1.1 × 10^6^ cells/well on fixed dermal equivalents, and they were cultured in growth medium under submersion conditions for 4 d in a humidified atmosphere at 37 °C with 5% CO_2_ and 12.5% O_2_. After maintenance in a submersion culture, the HSEs were exposed to the air-liquid interface in a humidified atmosphere at 37 °C with 5% CO_2_. The growth medium for culturing HSEs consisted of DMEM supplemented with 10% FBS, 1% penicillin/streptomycin, 5 µg ml^-1^ insulin (FUJIFILM Wako Pure Chemical Corporation), 500 < M ascorbic acid (FUJIFILM Wako Pure Chemical Corporation), 10 ng ml^-1^ bFGF (PeproTech, NJ, USA), 1 µM hydrocortisone (FUJIFILM Wako Pure Chemical Corporation), and 5 nM VEGF 165 Human Recombinant (PeproTech) and was replaced with fresh medium every 48 to -72 h.

### H&E staining and immunohistochemistry

For histological analysis, skin equivalents from three independent experiments were fixed with 4% formaldehyde and embedded in paraffin or Tissue-Tek O.C.T. compound (Sakura Finetek Japan Co., Ltd., Tokyo, Japan). Hematoxylin-eosin staining was performed on paraffin sections (10 µm thick). The stained sections were observed via AxioScan.Z1 (Carl Zeiss, Oberkochen, Germany). For fluorescence immunohistochemistry, frozen sections (10 and 50 µm) and paraffin sections (10 µm) were prepared and stained as previously described^23^. Details of the primary and secondary antibodies and associated epitope recovery methods are included in Table 1. All fluorescence microscopy images were captured with an AxioScan.Z1 or LSM 900 confocal microscope (Carl Zeiss).

**Table 1.**
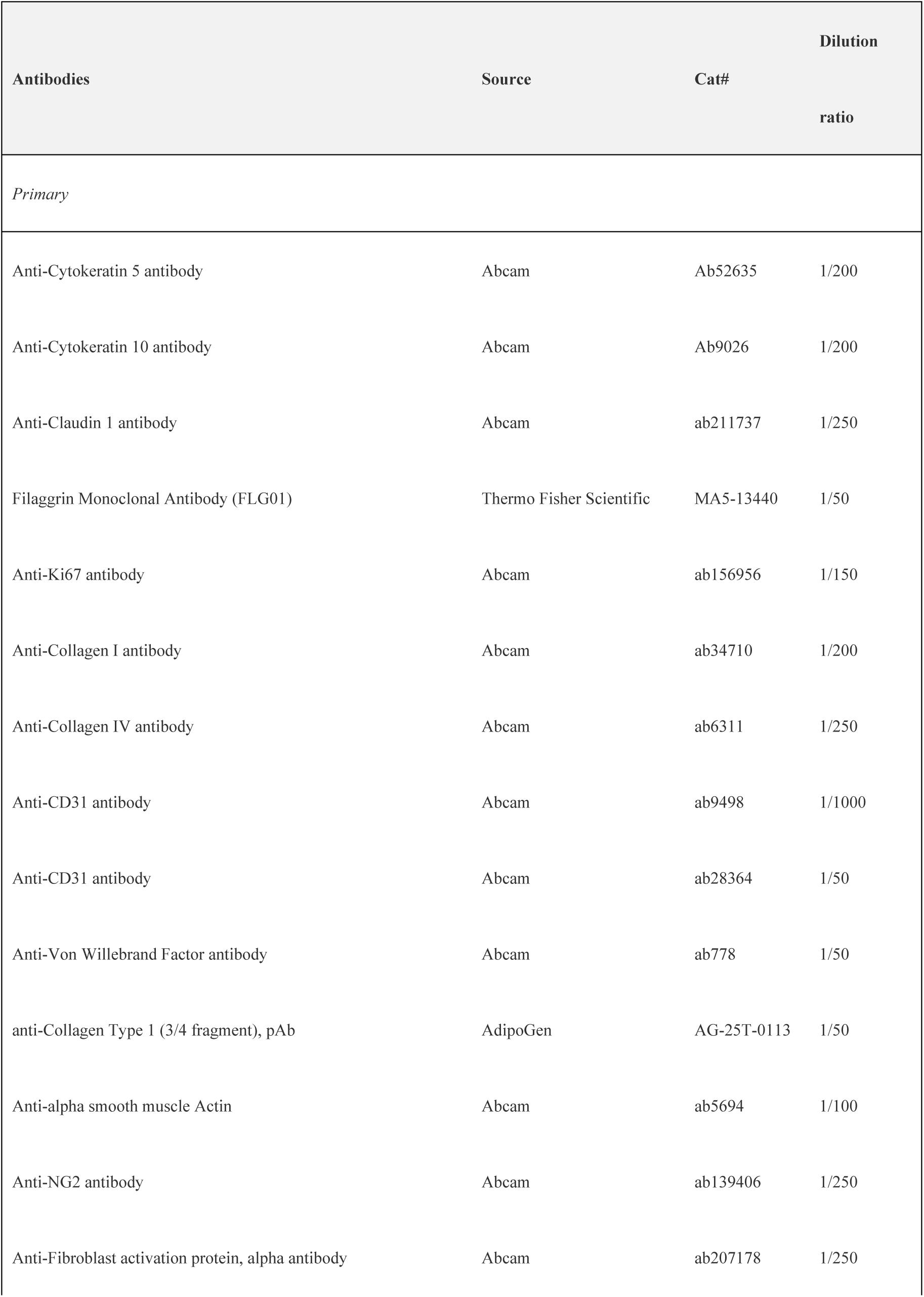
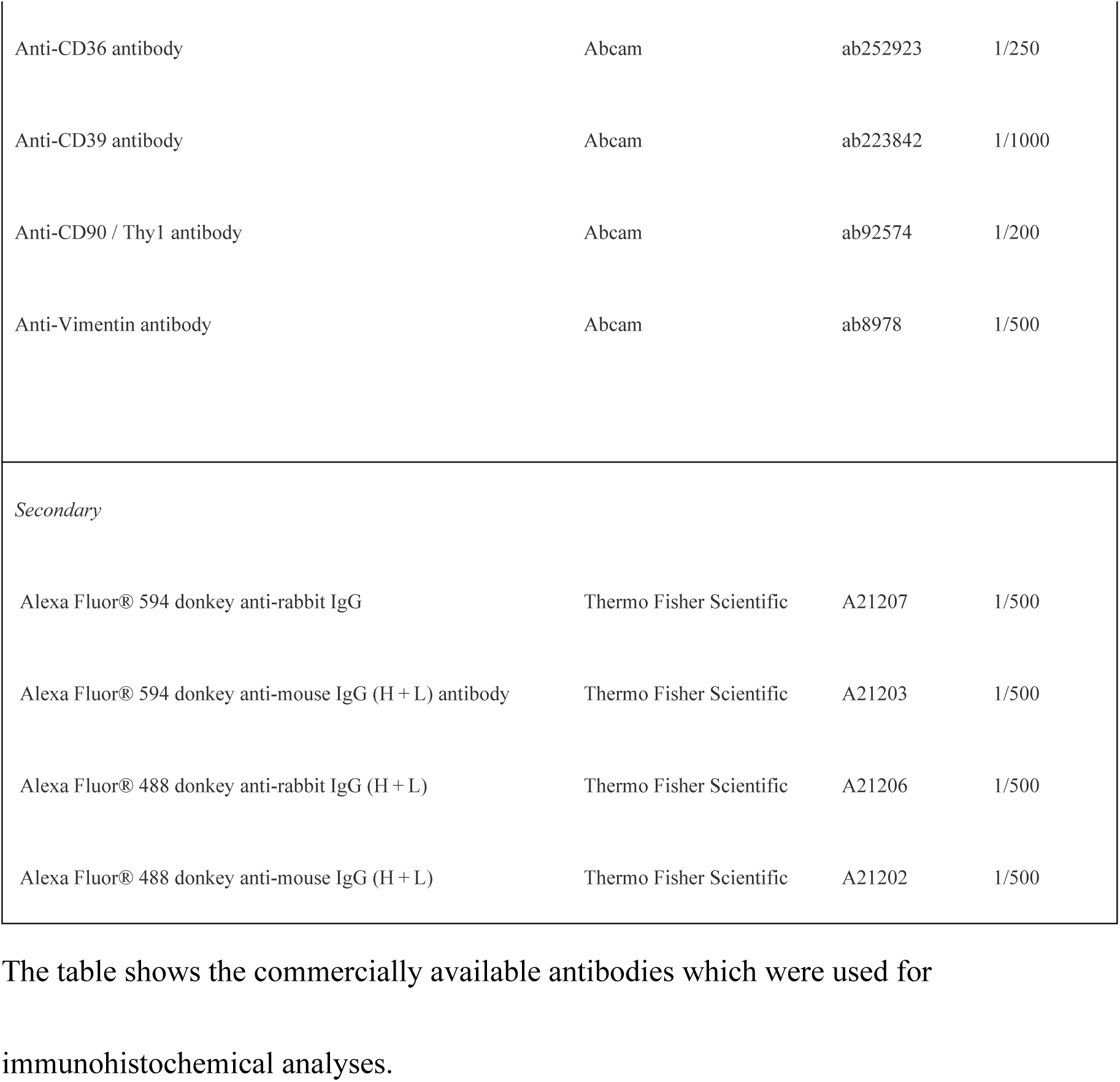
Primary and secondary antibodies.

### Quantitative analysis of epidermal cell proliferation

The proliferation rate of epidermal cells was quantified by counting keratinocytes with Ki67-positive nuclei located at the basal or immediately suprabasal epidermis. The measurement area was a total of 3 images obtained by randomly acquiring three z-stack images of 1000 × 340 × 10 μm from HSE tissues that were derived from three different wells.

### Quantification of blood vessel alignment

Anisotropic alignment of endothelial tubes and collagen fibers was assessed via two-dimensional fast Fourier transform (2D-FFT) analysis, as previously described^61^. Briefly, uncompressed images of endothelial tubes labeled with anti-CD31 antibody and basement membranes labeled with anti-COL1 antibody were analyzed with the FFT function of ImageJ, and the radial summation of the FFT frequency plot was calculated via the Oval profile plug-in. The degree of fiber alignment was reflected by the shape and height of the major peak in the FFT alignment plot. Images with oriented blood vessels resulted in a prominent peak centered at the principal axis of fiber alignment, whereas images with unaligned actin fibers resulted in an alignment plot with a broad peak or no peak.

### Single-cell RNA-seq library preparation

NHEKs, NHDFs, and HUVECs cultured to a subconfluent state were harvested through trypsinization according to the product protocol. Briefly, the cells were washed with HEPES-buffered solution (Kurabo) to remove the medium and treated with 0.025% trypsin/EDTA solution (Kurabo) for 3–7 min, followed by the addition of a trypsin-neutralizing solution (Kurabo) and centrifugation to obtain a cell pellet. Constituent HSE cells were harvested through enzymatic digestion. On day 14 of air-liquid interface culture, the epidermis and dermis of the HSE were mechanically separated using forceps. The epidermal layer was enzymatically dissociated with 0.025% trypsin/EDTA solution, while the dermal layer was incubated in 10% FBS-supplemented DMEM containing 5000 U/ml collagenase type 1 (Worthington Biochemical Corp., NJ, USA) in a 37 °C water bath. The resulting cell suspension was filtered through a 70-µm cell strainer to remove debris and centrifuged to obtain a cell pellet. Single-cell RNA sequencing was conducted using the Chromium Next GEM Single Cell 3ʹ Kit v3.1, Chromium Next GEM Chip G Single Cell Kit, and Dual Index Kit TT Set A (10x Genomics) according to the manufacturer’s protocol (CG000315 Rev E). Cell viability and count were assessed using a Countess 3 Automated Cell Counter following trypan blue staining and visual inspection. The cell suspension was processed on a chromium controller for gel bead-in-emulsion (GEM) generation, where polyadenylated mRNA was reverse transcribed to cDNA and uniquely barcoded. The cDNA was subjected to amplification, quality assessment via electrophoresis, and library preparation involving fragmentation, end-repair, A-tailing, adaptor ligation, and PCR amplification. Library quality was evaluated using Agilent’s High Sensitivity D5000 ScreenTape system, and the DNA concentration was measured with a Qubit 1X dsDNA HS Assay Kit on a Qubit 4 Fluorometer. Libraries were sequenced on an Illumina NovaSeq 6000 with PE150, generating approximately 120 Gb of data per sample from ∼800 million reads.

### Single-cell RNA-seq data processing: *In vitro* skin data

FASTQ reads of 10× scRNA sequencing data were processed with the hg38 reference genome via Cell Ranger (v7.1.0)^62^. A total of 27,082 cells passed the quality control steps. The processed matrices of different batches were merged, and the following analyses were performed via SEURAT-1 (v5.0.3). Cells with 1,000 to 8,000 RNA features and mitochondrial RNA contents of less than 7% were selected. The selected expression matrix was normalized by the total number of UMIs per cell and was log-transformed.

The filtered dataset was split into original sample identities via the Split Object function, followed by data log normalization of UMI counts and identification and the identification of 2000 more variable genes per sample.

The integrated data were then used for standard cell clustering and visualization with Seurat, which uses the 2000 most variable genes of the integrated dataset as input. Next, we used the function FindIntegrationAnchors with default parameters and 30 canonical correlation analysis (CCA) dimensions to identify the integration anchors between our five datasets. These anchors were subsequently used for integration via the IntegrateData function, again with the first 30 CCA dimensions and default parameters.

The integrated data were then used for standard cell clustering and visualization with Seurat, which uses the 2000 most variable genes of the integrated dataset as input. First, the data were scaled via the ScaleData function, and principal component analysis (PCA) dimensions were calculated via the RunPCA function. Next, unsupervised clustering of the data was performed with the FindNeighbors and FindClusters functions. For the FindNeighbors function, we used the first 30 PCA dimensions to construct a shared nearest neighbor (SNN) graph for our dataset. Then, we clustered the cells with the function FindClusters via a shared nearest neighbor (SNN) modularity optimization-based clustering algorithm with a resolution of 0.7. Finally, for visualization, we used the RunUMAP function with default parameters and 30 PCA dimensions.

### Single-cell RNA-seq data processing: *In vivo* skin data and data integration with *in vitro* skin data

*In vivo* skin single-cell analysis was carried out via methods described in previous studies^11^. After mapping cell types based on the expression of marker genes for each cell type, we further explored the correspondence between the clusters and the four fibroblast subpopulations mentioned in the original paper. This was accomplished by examining the expression of marker genes for papillary fibroblasts and reticular fibroblasts, as well as the top enriched Gene Ontology (GO) terms for the four fibroblast clusters. Consistent with previous studies, Cluster 1 was classified as secretory-reticular, Cluster 2 as proinflammatory, Cluster 3 as secretory-papillary, and Cluster 9 as mesenchymal.

To integrate *in vitro* and *in vivo* data, cells annotated as vascular ECs, pericytes, keratinocytes, and fibroblasts were initially extracted from the *in vivo* dataset. The cells from each sample were filtered under specific conditions: for *in vitro* samples, cells with 1000 to -8000 RNA features and mitochondrial RNA contents of less than 7% were selected; for *in vivo* samples, the criteria were fewer than 7500 RNA features and mitochondrial RNA contents of less than 5%. Following this, normalization was performed via the SCTransform method. Using Seurat’s default approach, the top 2000 genes with the most variability from each dataset were extracted. All the data were then combined to compute principal component analysis (PCA) dimensions, and batch effects between datasets were mitigated using the Harmony algorithm’s RunHarmony unction (parameters set to theta=1, max.iter.harmony=10). Unsupervised clustering of the data was subsequently performed via the FindNeighbors and FindClusters functions. Finally, for visualization purposes, the RunUMAP function was employed with default parameters and 30 PCA dimensions.

### Cell–to-cell communication analysis

To analyze the estimated intercellular interactions established by different cell types present in artificial skin, we utilized R CellChat3 (v1.6.1)^63^. This tool can quantitatively infer and analyze intercellular communication networks from scRNA-seq data and contains ligand-receptor interaction databases (http://www.cellchat.org/). In our approach, pairwise comparisons across all cell clusters present in the artificial skin dataset were conducted, and the analysis was performed using the default parameters set by the tool.

### Evaluation of the epidermal barrier function of HSEs via TEWL measurement

Trans-epidermal water loss (TEWL) was measured in the HSEs (*n* = 4) using a Tewameter TM HEX (Courage & Khazaka electronic, Cologne, Germany) according to the instruction manual. Prior to collection, the skin equivalent was incubated inside the biosafety cabinet for 15 min to allow equilibration to the ambient temperature and humidity of the room. The measurement was repeated three times, and the average and standard deviation of each well were calculated.

### Measurements of skin elasticity

Skin elasticity was determined via a noninvasive, *in vivo* suction skin elasticity meter, the Cutometer MPA 580 (Courage & Khazaka electronic). The measurement conditions were the same as those used in previously reported human clinical studies^41^. Briefly, with a 2-mm probe, a negative pressure of 400 mbar was applied to the HSEs for a period of 2 s, followed by 2 s of relaxation time, and the ratio of immediate retraction (Ur) and the ability of redeformation of the skin (Ua) to final distension (Uf) was analyzed (R2 = Ua/Uf, R7 = Ur/Uf).

### Nutrient-poor culture of HSEs and evaluation of their responsiveness to ascorbic acid

All HSEs were treated with nutrient-poor medium (DMEM supplemented with 1% FBS, 1% penicillin/streptomycin, 5 µg ml^-1^ insulin, and 1 µM hydrocortisone) at the air-liquid interface on day 3 and maintained for 12 d. Then, 500 μM ascorbic acid was added to the HSEs cultured at the air-liquid interface on day 5 and maintained for 10 d.

### Statistics and reproducibility

Microsoft Excel (Microsoft, Redmond, WA, USA) and BellCurve for Excel were used for statistical analysis. All values are expressed as the means ± standard deviations (SDs). Analysis of the samples was performed at least in triplicate, and the results were averaged. Differences between groups were considered significant at P<0.05. All the experiments were repeated at least two times.

## Data availability

scRNA-seq data from this study have been deposited into the Sequence Read Archive (SRA) database under the BioProject ID. PRJNA1191968. All data are available from the corresponding authors upon reasonable request.

## Acknowledgments

We thank laboratory members at the Institute of Advanced Biomedical Engineering and Science, TWIns, Tokyo Women’s Medical University. We also thank Mr. M. Hashimoto, Y. Hayashi, and all the members of Rohto Pharmaceutical Co., Ltd., for their support and encouragement in this project. We would like to thank Rohto Pharmaceutical Co., Ltd., for funding support.

## Author contributions

S.K., S.S. and T.S. designed the research plans; S.K. and S.Y. performed the experiments; S.K., S.S., T.K., S.Y., M.H., and T.S. discussed the results; and S.K. and S.Y. wrote the manuscript.

## Author information

Correspondence and requests for materials should be addressed to S.K.

## Supplementary Information

**Supplementary Figure 1.**
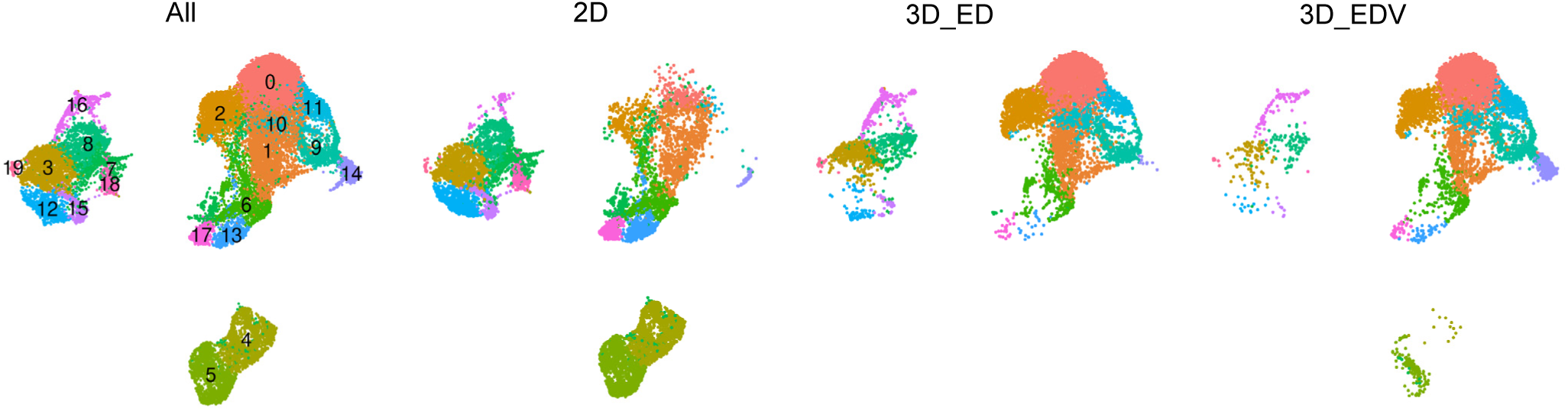
scRNA-seq analysis of all *in vitro* samples. UMAP plot of scRNA-seq for *in vitro* human skin data, and distribution of all cells under each of the three conditions (2D, 3D_ED, and 3D_EDV).

**Supplementary Figure 2.**
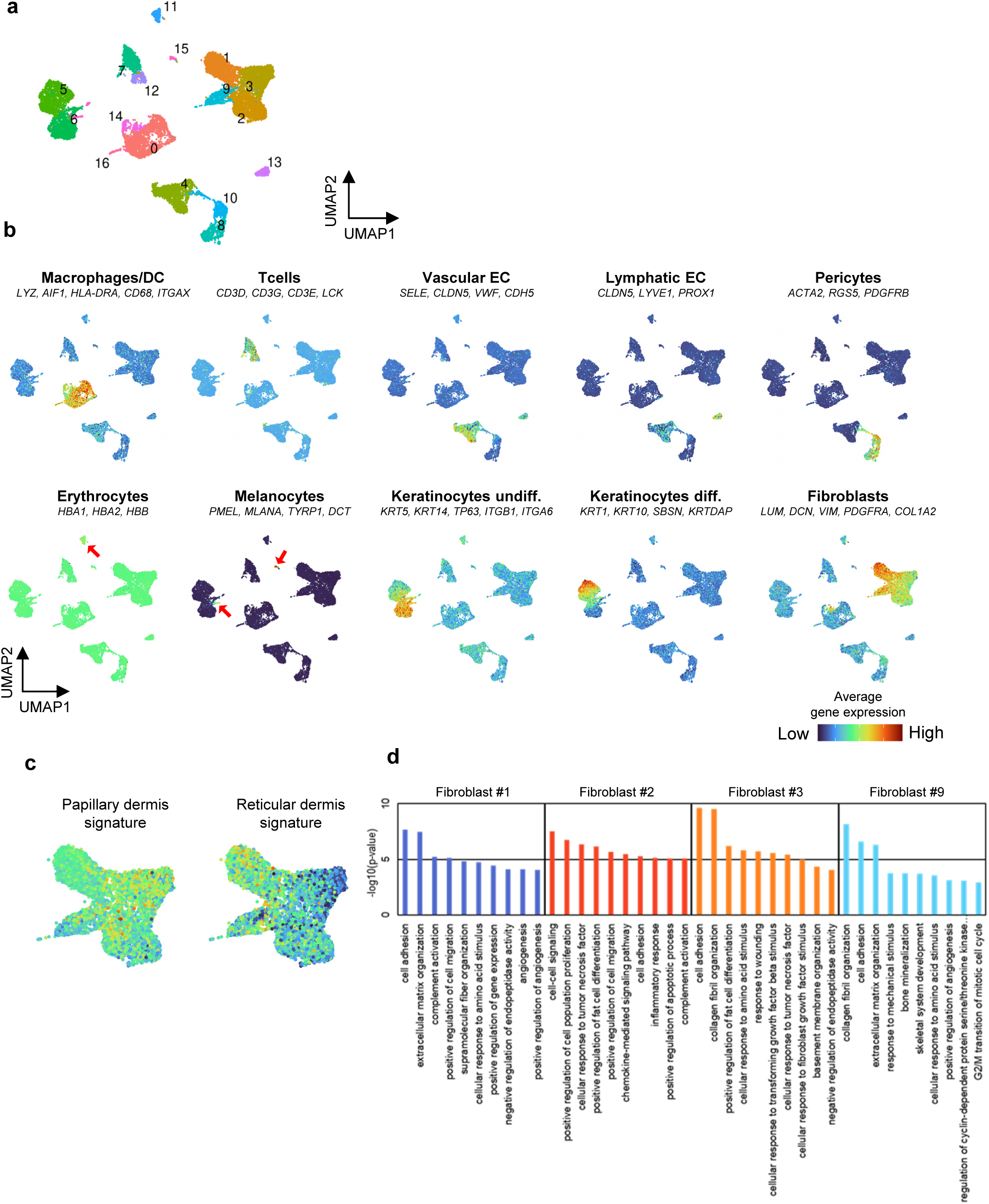
Re-evaluation of *in vivo* samples through scRNA-seq. (a) A UMAP plot presenting single-cell transcriptomic data from whole human skin samples (n = 5). Each point signifies a single cell, with coloration based on unsupervised clustering executed using Seurat. (b) Mean expression of genes forming the papillary and reticular gene signatures applied to predict the dermal localization of fibroblasts within the four clusters. (c) The 10 top significantly enriched GO terms within each fibroblast subpopulation, arranged by *P*-value (d) A UMAP plot featuring the average expression of previously established cell type markers for distinguishing cell populations. Red denotes maximum gene expression, whereas blue represents minimal or non-existent expression of a specific gene set in log-normalized UMI counts.

